# Predictors of taxonomic splitting and its role in primate conservation

**DOI:** 10.1101/2021.09.10.459781

**Authors:** Maria J.A. Creighton, Alice Q. Luo, Simon M. Reader, Arne Ø. Mooers

## Abstract

Species are the main unit used to measure biodiversity, but different preferred diagnostic criteria can lead to very different delineations. For instance, named primate species have more than doubled in number since 1982. Such increases have been attributed to a shift away from the ‘biological species concept’ (BSC) in favour of less inclusive species criteria. Critics of recent changes in primate taxonomy have suggested taxonomic splitting may be biased toward certain clades and have unfavourable consequences for conservation. Here, we explore predictors of taxonomic splitting across primate taxa since the initial shift away from the BSC nearly 40 years ago. We do not find evidence that diversification rate, the rate of lineage formation over evolutionary time, is significantly linked to splitting, contrary to expectations if new species concepts and taxonomic methods identify incipient species. We also do not find evidence that research effort in fields where work has been suggested to motivate splitting is associated with increases in species numbers among genera. To test the suggestion that splitting groups is likely to increase their perceived risk of extinction, we test whether genera that have undergone more splitting have also observed a greater increase in their proportion of threatened species since the initial shift away from traditional taxonomic methods. We find no cohesive signal of taxonomic splitting leading to higher threat probabilities across primate genera. Overall, this analysis sends a positive message: threat statuses of primate species are not being overwhelmingly affected by splitting. Regardless, we echo warnings that it is unwise for conservation to be reliant on taxonomic stability. Species (however defined) are not independent from one another, thus, monitoring and managing them as such may not meet the overarching goal of conserving biodiversity.

## INTRODUCTION

“Species” are an integral unit of biodiversity used across many sub-disciplines of biology, yet how scientists define species has been subject to change. Notably, in the last 40 years, the emergence of new methods for identifying diagnostic differences between populations (e.g., molecular phylogenetic methods) and changes in preferred species criteria have led to large increases in species numbers across many clades (Agapow *et al*., 2004).

Groves (2014) provides a brief overview of popular species definitions employed by taxonomists through the late nineteenth to twentieth century. One notable phenomenon is the considerable decrease in diagnosed species that occurred following the rapid adoption of the polytypic species concept beginning in the 1890’s. The polytypic species concept emphasizes that species should be inclusive and that one should delineate taxa that resemble one another as subspecies (Groves, 2014). The popularization of the polytypic species concept was eventually accompanied by the widespread adoption of the ‘biological species concept’ (BSC) beginning in the early 1960’s (Groves, 2014). The BSC defines species as populations/meta-populations that do not interbreed with other populations/meta-populations under natural conditions (Mayr, 1963; Groves, 2014). While this definition has been subject to revisions, the central premise of the BSC is that reproductive barriers are key to diagnosing species (Groves, 2014). The BSC was widely accepted and layered onto the pre-existing polytypic species concept, creating a period of relative taxonomic stability for vertebrates from the 1960’s to 1980’s (Isaac *et al*., 2004). However, various criticisms of the BSC did emerge, the most notable being the practical difficulty of diagnosing species under the BSC because of the need for information on reproductive barriers (Donoghue, 1985; Tattersall, 2007). Such criticisms suggested the need for new approaches to delineating species that offered higher diagnostic power.

In the last 40 years the ‘phylogenetic’ or ‘diagnostic’ species concept (PSC) (Cracraft, 1983) has been widely popularized in vertebrate taxonomy due to its diagnosable advantages over previous species definitions (Isaac *et al*., 2004; Cotton *et al*., 2016). Under the PSC, a species is diagnosed as the smallest population or meta-population that is distinct in heritable differences from other populations or meta-populations (Cracraft, 1983; Groves & Grubb, 2011; Groves, 2014). According to its proponents, the PSC’s emphasis on diagnosable evidence gives it an advantage over other species concepts because it allows users to rely on a range of data types, including newly available molecular markers, to make distinctions (see, e.g., the variety of data types used to describe the newest species of ape, *Pongo tapanuliensis*; Nater *et al*., 2017). Together, new species concepts and methodological advancements have characterized a shift away from biological species and toward species that are delineated based on distinctive, diagnosable differences.

Although using distinctive differences to delineate species offers advantages, this approach has also been subject to criticism, notably, for its tendency to split species into a range of less-inclusive units (Agapow *et al*., 2004; Zachos *et al*., 2013; Zachos & Lovari, 2013). Many populations previously recognized as subspecies or morphological variants have been elevated to the full species status, resulting in a large increase in the number of listed species. For instance, 181 species of primates were listed by Honacki *et al*. (1982), one year prior to Cracraft’s (1983) proposal of the PSC. Today the IUCN (International Union for Conservation of Nature) lists over 500 distinct primate species (Estrada *et al*., 2017). Some families (e.g., Cheirogaleidae and Indriidae) have more than tripled in species richness over the past ∼40 years (see Figure 1). While some new species have been added as a result of new field discoveries, a majority are populations which were previously identified at lower taxonomic levels that have now been elevated to species status following a shift in the accepted approach to diagnosing primate species (Tattersall, 2007). Some attribute this taxonomic increase to the popularity of the PSC (see, e.g., Isaac *et al*., 2004), while others credit increased exploration and the development of new techniques for evaluating diagnosability (see, e.g., Köhler *et al*., 2005; Harris & Froufe, 2005; Padial & De la Riva, 2006; Sangster, 2009). The trend of increasing species numbers by raising taxonomic statuses has been referred to as ‘taxonomic inflation’ by some (Isaac *et al*., 2004; Rylands & Mittermeier, 2014) and has been criticized for being non-random, and biased toward certain clades (Isaac *et al*., 2004; Agapow *et al*., 2004; Zachos *et al*., 2013; Zachos & Lovari, 2013; Rylands & Mittermeier, 2014).

**Figure 1:**
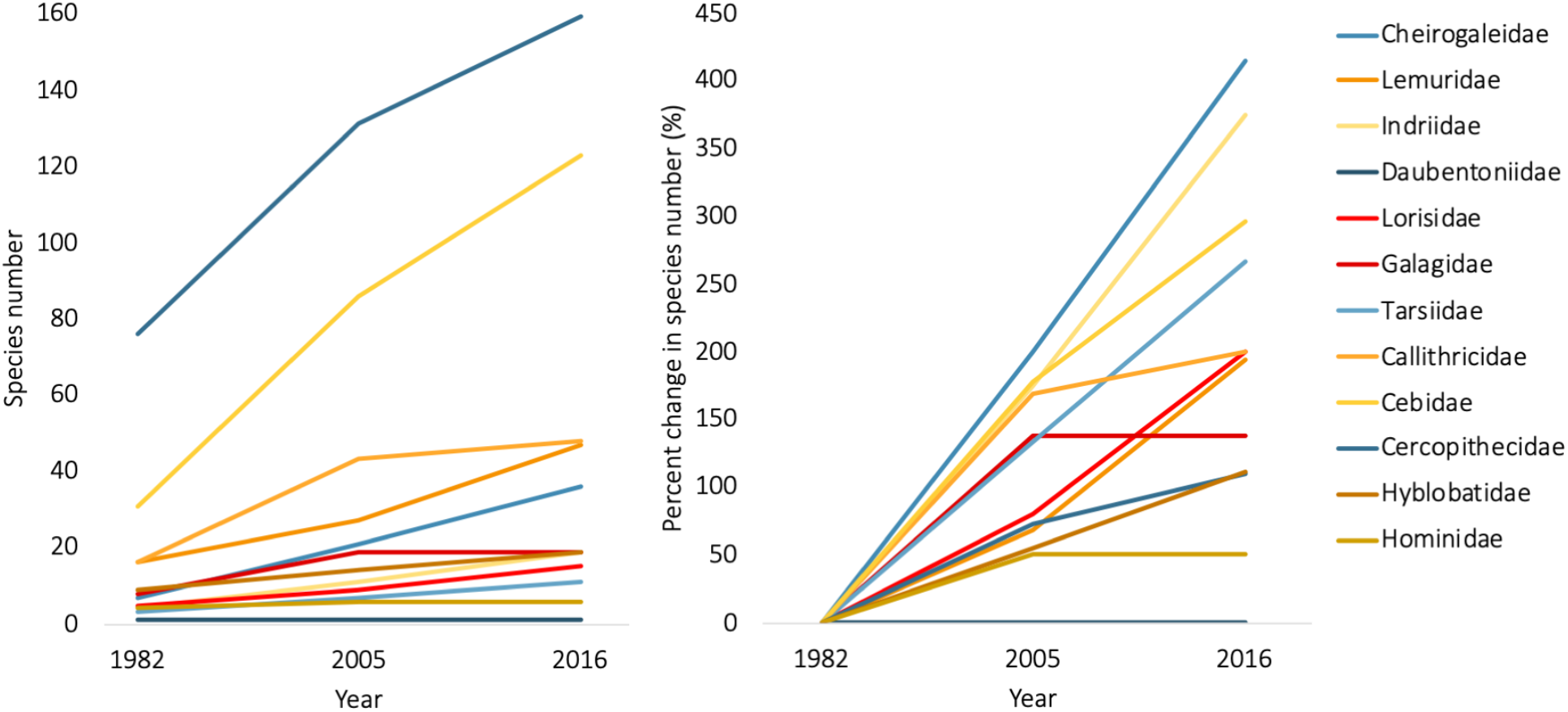
Species numbers and percentage change in species numbers for historic primate families recorded at three time points by Honacki *et al*. (1982), Wilson & Reeder (2005) and the IUCN species list from 2016 (data documented in Estrada *et al*., 2017).

Disproportional splitting among taxonomic groups could have several causes. Variation in the number of new species described across taxa could be driven by variation in the rate at which lineages evolve, such that new species descriptions are tracking cryptic diversity or incipient species formation among rapidly evolving taxa. In this case, new species listings might point to situations where there is a discordance between patterns of genetic change and the evolution of gross morphological changes used by traditional taxonomists. This could be due to ecology – if some lineages are diversifying along ecological axes not captured in traditional taxonomic approaches – or due to demographics – if some lineages have ecologies and/or histories that lead to faster local genetic coalescent times and so diagnosability. Under either of these scenarios, variation in which clades observe the greatest amount of splitting under new taxonomic approaches would simply reflect underlying biological reality. However, it is also possible that splitting is driven by other factors that may be prone to bias. Critics have argued that many increases in species numbers are artificial, reflecting major shortcomings of new species criteria and a reliance on insufficient data (Zachos *et al*., 2013; Zachos & Lovari, 2013). Zachos *et al*. (2013) provides evidence suggesting unwarranted splitting in select cases, advocating that splitting has been taken to a “molecular extreme”. Many different types of molecular data are used to justify splitting (e.g., genetic data from mitochondrial DNA barcoding) and, by this logic, groups may continue to be split as an increasing amount of molecular data become available for them. If true, this would lead to the continual identification of increasingly exclusive diagnostic features such that we are likely to find new species the more we look for them.

It is also possible that conservation interest in particular groups could motivate splitting. Limited funding for conservation research is increasingly focused on ‘biodiversity hotspots’ and it has been suggested that researchers could have a vested interest in declaring taxa in these regions as endemic species (Karl & Bowen, 1999; Isaac *et al*., 2004). There is evidence from some charismatic groups that taxa receive more conservation attention and funding when comprised of multiple, small, and taxonomically distinct populations (e.g., African apes; Stanford, 2001; Oates, 2006; Gippoliti & Amori, 2007). Species lists are often used to determine which groups should receive conservation attention (Mace, 2004) and so changing the way we define species may also change which groups receive action.

Along with being potentially biased toward certain taxonomic groups and related to the point made above, taxonomic splitting has also been suggested to result in individually more imperiled populations which could distort conservation agendas (Agapow *et al*., 2004; Isaac *et al*., 2004; Morrison *et al*., 2009; Zachos, 2015; Robuchon *et al*., 2019). One criterion used by the IUCN to classify species as imperiled is population size: species may be designated “Vulnerable” if there are fewer than 1000 mature individuals found in the wild and “Endangered” if there are fewer than 250 (Agapow *et al*., 2004; Frankham *et al*., 2012). Thus, splitting one species into several new species may result in one or more receiving a (more) imperiled status (Agapow *et al*., 2004; Isaac *et al*., 2007). This could lead to seemingly rare but poorly defined species being prioritized over well-defined and perhaps biologically more distinctive species (Pillon & Chase, 2007). Recent evidence suggests species splitting is not a driver of threat status for birds (Simkins *et al*., 2020), however, it is unknown whether these findings are generalizable across other taxonomic groups (Garnett & Thomson, 2020).

Here, we set out to better understand the causes and consequences of taxonomic splitting in primates. We test (i) predictors of taxonomic splitting, that is, whether recent taxonomic increases are associated with the amount of research being done in fields suggested to motivate splitting or with a lineage’s underlying diversification rate, and (ii) impacts of taxonomic splitting, that is, whether rates of splitting dictate which groups are most imperiled. Because some newly described primate species have previously been described as subspecies or subpopulations of more than one different biological species prior to being assigned to the full species rank (which would make quantifying rates of splitting at the species-level difficult), we ask these questions at the genus level.

To explore our first question regarding predictors of taxonomic splitting, we consider whether biological factors or measures of human-induced bias explain increases in species numbers. To test potential human-induced biases in splitting, we consider broad estimates of research effort for each taxon and predict that more research done on a given taxon may be associated with more splitting. We examine research effort in two fields: molecular genetics (since molecular work could cause species to be split continuously as finer molecular distinctions are made) and conservation (since splitting has been suggested to be motivated by conservation interests). To explore possible biological explanations for trends in taxonomic splitting across clades, we test whether recent taxonomic increases are explained by diversification rate. Clades with high recent diversification rates are expected to contain more incipient or cryptic species than lineages diversifying at a lower rate since they will contain more closely related lineages that resemble one another. Therefore, if new approaches help to identify incipient or cryptic species, diversification rate should be positively correlated with splitting.

To explore our second question linking splitting and risk, we test whether increases in species numbers are associated with a change in the number of threatened species listed in genera through time. Ideally we would look at changes in a weighted measure of threat score that differentiates between threat categories of varying severity (e.g., using the Red List Index (RLI); Butchart *et al*., 2007; Bubb *et al*., 2009); however, criteria for inclusion in Red List threat categories have changed considerably over time, meaning weights assigned to categories for RLI calculations do not match up with categories used in the past (see, e.g., statuses in IUCN Conservation Monitoring Centre, 1986). Because of these changes in Red List criteria, we instead ask whether clades with species that have been split more frequently have observed a greater increase in their proportion of threatened species (defined below) through time in comparison to those which have been split less frequently. We predict that if splitting is driving an increase in threatened species, we should observe a positive association between taxonomic increases caused by splitting and change in the proportion of threatened primate species.

## MATERIALS AND METHODS

### Data

To measure the number of primate species that were described before the introduction of the PSC and new molecular techniques, we used the last pre-PSC taxonomy, that of Honacki *et al*. (1982). This taxonomy contains 181 species and is considered a reliable estimate of the number of species thought to exist during the popularity of the BSC (see Rylands & Mittermeier, 2014). We then recorded if each species in this taxonomy was historically considered to be threatened by referencing the most complete IUCN Red List published around the same time (IUCN Conservation Monitoring Centre, 1986). Honacki *et al*. (1982) was contrasted with the IUCN taxonomy from 2016 and attendant data documented in Estrada *et al*. (2017), which lists 503 species. For each of these 503 species we noted their taxonomic placement (genus and family), whether or not they were considered threatened (VU=Vulnerable, EN=Endangered, or CR=Critically Endangered), and their biogeographic region. We note that while IUCN assessments have some shortcomings (see, e.g., Rueda-Cediel *et al*., 2018), the IUCN provides the largest global data on threat status and is influential in determining how most species are managed. For each species described by the IUCN in 2016 that was not listed in Honacki *et al*. (1982), we scored whether the species was a “*de novo*” species description (Burgin *et al*., 2018): there were 40 such cases where a new species had not been previously formally identified as a subspecies or subpopulation of another species prior to splitting. These cases represented new species descriptions where it was unclear whether a new species was a result of taxonomic splitting or the discovery of an entirely new population.

We compiled all the species listed by the IUCN in 2016 into 12 families and 50 genera found in Honacki *et al*. (1982). Family name “Callimiconidae” in Honacki *et al*. (1982) was not used as this taxon has since been recognized as a genus of the larger family “Callitrichidae” (Wilson & Reeder, 2005). We removed *Rungwecebus kipunji* from the IUCN species list from 2016 as it represents a newly discovered genus that does not collapse into any of the genera provided by Honacki *et al*. (1982).

Research effort in the fields of molecular genetics and conservation was estimated for each genus through a literature review of publications in the Web of Science Core Collection published between 1983 (when a general trend toward taxonomic splitting first began) and 2016. All search terms for these literature reviews and further details on related methodology are documented in the supplementary materials (Tables S1 and S2). In this study we used the genera listed in Honacki *et al*. (1982) (n=50), many of which have since been further separated into multiple genera. Thus, when appropriate we included new genus names in addition to those listed by Honacki *et al*. (1982) in the literature searches (see Table S3). In total, our final sample was 688 publications on molecular genetics and 2222 on conservation.

Diversification rate was estimated with the method-of-moments approach described in Magallon & Sanderson (2001) (i.e., ln(taxa richness)/stem age). Diversification rate estimates generated using this method often rely on species numbers as their estimate of taxa richness, meaning that diversification rate estimates are inherently biased by the splitting phenomenon we are studying (i.e., frequently split genera will receive disproportionally high diversification rates). Therefore, richness scores for our diversification rate calculations were determined as the counts of the well-resolved “lineages” described in Creighton *et al*. (2021). These lineages were determined by creating a time cut-off in the 10kTrees consensus primate phylogeny (Arnold *et al*., 2010) in an attempt to eliminate very young newly described species and obtain a consistent (unbiased) estimate of diversity across clades. These lineages were assigned to each of the 50 genera described in Honacki *et al*. (1982). Diversification rate was then estimated by taking the natural log lineage richness for each genus and dividing by the stem age of that genus (Magallon & Sanderson, 2001). Stem ages for each genus were extracted from the 10kTrees consensus phylogeny (version 3) (Arnold *et al*., 2010) trimmed to contain a single branch representing each genus. During this process, there were several instances where genera described in Honacki *et al*. (1982) were non-monophyletic within the more recent primate phylogeny we used (Arnold *et al*., 2010), making it unclear how to assign a divergence date for these clades. We therefore removed eight genera from the analyses where diversification rate was a variable of interest: *Presbytis, Lemur, Galago, Cebuella, Cercocebus, Cercopithecus, Papio*, and *Pygathrix*.

### Analysis

To test our questions about the predictors of species-splitting and its consequences for conservation, we fit a series of linear effects and mixed effects models. We note that the response variables used in these analyses (i.e., measures of taxonomic increase and extinction risk) are likely to be phylogenetically clustered, and phylogenetic models could be used to account for this influence of phylogeny; however, many genera listed by Honacki *et al*. (1982) are non-monophyletic, making it unclear how to designate them a single branch in modern phylogenies (see discussion above on diversification rate). Importantly, after accounting for regional differences, family contributed little to no variance in any of our models, indicating that phylogenetic relationships at that level were not confounding our results. We provide further discussion on model choice in the supplementary materials.

Data were analysed using R version 4.0.5 (R Core Team, 2021).

#### Predictors of Taxonomic Splitting

To determine if measures of potential human bias (i.e., research effort) or diversification rate explained discrepancies in splitting across taxa, we tested to see if these variables were significantly associated with the number of species added to primate genera since 1982 while controlling for the original number of described species (i.e., biological species) and region. We fit three generalized linear mixed effects models with Poisson distributions using the lme4 package in R (Bates *et al*., 2014), and obtained p-values using the lmerTest package (Kuznetsova *et al*., 2017). In these models the response variable was the number of species assigned to a given genus by the IUCN in 2016 that had not been previously described by Honacki *et al*. (1982). Each model had either conservation research effort, molecular genetics research effort or diversification rate included as a fixed effect, as well as biogeographical region and the number of species in the genus per Honacki *et al*. (1982) to control for their effects on splitting. The natural logarithm (ln) of the number of species listed for each genus in Honacki *et al*. (1982) was included as both a linear and a quadratic term following inspection of raw plots and plots against scaled residuals from the simulation output. A square-root transformation was used on molecular genetics research effort to decrease the impact of outliers on model fit. To assist with model stability and convergence, we scaled all continuous variables in the model to have a mean of zero and standard deviation of one (Becker *et al*., 1988). Mainland Africa and Asia (hereafter mainland Africa + Asia) were grouped together and served as the baseline region in our models based on previous studies that have shown that the taxonomy of primates from these regions has been relatively stable compared to Madagascar and the Neotropics (e.g., Isaac & Purvis, 2004; Isaac *et al*., 2004; Tattersall, 2007). We also chose to group Asia and Africa together because one genus (*Macaca*) is found in both regions. Genus ID was included as a random effect in these models to correct for overdispersion. Family (nested within region) was originally included as a random effect but contributed very little to model fit and created issues with convergence due to overfitting, and thus was dropped from the final models. We tested potential interaction terms with all variables and region to test for regional effects, but none were significant and so these terms were also dropped from the final models. Before running models with other predictors included, we also ran a model including only the linear and quadratic terms for the number of species listed for each genus in Honacki *et al*. (1982) as predictors to assess their association with taxonomic increase in the simplest model. We ran all models a second time after removing the 40 *de novo* species from our response (Tables S4, S5 and S6). We checked model assumptions and fit by plotting residuals versus the fitted values and versus each covariate in the model. Residual plots and analyses done with the Diagnostics for Hierarchical Regression Models (DHARMa) R package (Hartig, 2017) indicated acceptable model fits.

#### Taxonomic Splitting and Threat Score

To test whether taxonomic splitting over time is associated with a change in the proportion of threatened species within genera, we conducted a two-step (hierarchical) analysis on species’ threat probability between 1982 to 2016. We first fit a generalized linear mixed effects model with a binomial distribution for the number of threatened and non-threatened species within a genus using the lme4 package in R (Bates *et al*., 2014). In this first model, the response estimated the proportional counts of species at risk within genera (equivalent to the per-species probability of threat) in the periods of 1982 and 2016. Predictors for this model included fixed effects for the time period (the baseline of 1982 and the change to 2016), the region that encompasses each genus’ natural distribution (Madagascar, Neotropics, mainland Africa + Asia), and an interaction between region and time period to account for geographic differences in threat probabilities through time. Genus identity was included as a random intercept, to account for repeated measures in 1982 and 2016, and as a random slope with time period to account for differences in changes to threat probabilities among genera. As discussed above, we included the taxonomic rank of family (nested within region) as a random effect but found that the variance in threat probabilities among families was minimal and that including this term also created convergence issues; given this, we subsequently omitted family from our models. Residual plots and analyses with the DHARMa R package (Hartig, 2017) indicated acceptable model fits for the final model. From this first model describing genus level changes in threat probabilities, we then used the REextract function implemented in the merTools package in R (Knowles & Frederick, 2020) to extract the genus level random slopes for time period and their associated standard errors, giving us an estimate of the varying effect of change in threat probability among genera (while conditioning on regional trends) between 1982 and 2016. We then fit a linear model where varying effect of change in threat probability from 1982 to 2016 was the response, and the proportional change in species within each genus (i.e., the number of new species in the IUCN species list from 2016 / original number of species in Honacki *et al*. (1982) – our measure of taxonomic increase) – was the predictor. We weighted each estimate of genus level change in threat probability by its standard error (w = 1/SE) to propagate the error of model estimated random effects. We ran this model a second time after removing *de novo* species descriptions from our response (Table S7). Raw data used for these tests are visualized in Figure 2.

**Figure 2:**
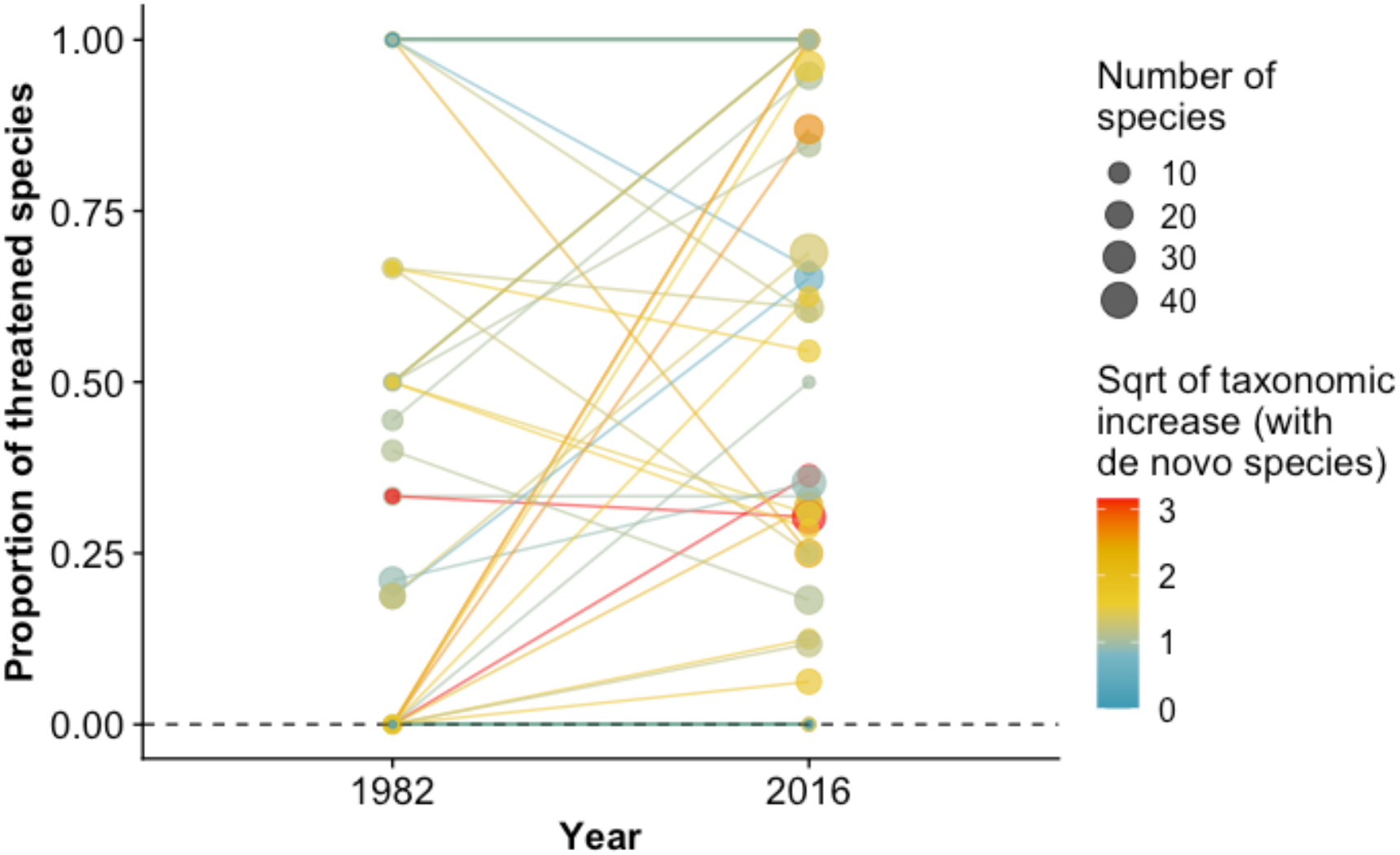
Scatterplots with trendlines showing the change in the proportion of species identified as being threatened in primate genera in 1982 and 2016 painted by the square root of taxonomic increase (including *de novo* species). Total number of species in each genus is indicated by point size.

Plots of the raw data indicated that patterns in taxonomic increases, the proportion of species at risk in primate genera today, and the changes in the proportion of threatened species in primate genera between 1982 and 2016 appeared to show regional differences (Figures S1 and S2). We thus ran a subsequent set of models to test for regional variation in the effect of splitting. In this analysis, we removed region from the first generalized linear mixed effects model and subsequently included region interacting with taxonomic increase as a predictor of the varying effect of change in genus threat probability in the second-order model. However, interaction effects were not significant in this model and thus we only considered the results of the first set of models reported above.

## RESULTS

### Predictors of Taxonomic Splitting

None of our measures – conservation research effort (β = -0.220; p = 0.180; Table S4), molecular genetics research effort (β= -0.188; p= 0.212; Table S5) or diversification rate (β= 0.132; p= 0.477; Table S6) – were significantly associated with increases in species numbers across primate genera. Removing *de novo* species did not impact this pattern of results (Tables S4, S5 and S6). Notably, in addition to sharing a significant linear relationship with taxonomic increase as expected (Tables S4, S5 and S6), the quadratic term added for the original number of species in 1982 was significant in the model with diversification rate, indicating a downwardly concave association with taxonomic increase regardless of whether de novo species were included (Table S6); this term had a p-value less than 0.1 in models that included conservation or molecular genetics research effort as predictors (Tables S4 and S5), indicating that the positive increase in splitting by initial genus size decelerates as genera become very large, even after accounting for other variables. We note that this quadratic term was significant without other predictors (i.e., research effort or diversification rate) present in the model, both with *de novo* species included in the response (β= -2.268; p= 0.011) and without *de novo* species included in the response (β= - 2.086; p= 0.013). In all three models without *de novo* species, taxonomic increases were significantly higher in the Neotropics compared to Africa and Asia (Tables S4 to S6).

### Taxonomic Splitting and Threat Score

Visual inspection of the raw data on taxonomic increases and proportion of species at risk for all genera revealed no clear indication that highly split genera have observed a disproportional increase in perceived extinction risk (Figures 2 and S3). Results from our linear model confirmed that taxonomic increases did not predict a change in threat score through time (β= 0.001; p= 0.975; Table S7; Figure 3). Removing *de novo* species did not impact this pattern of results (Table S7; Figure S4).

**Figure 3:**
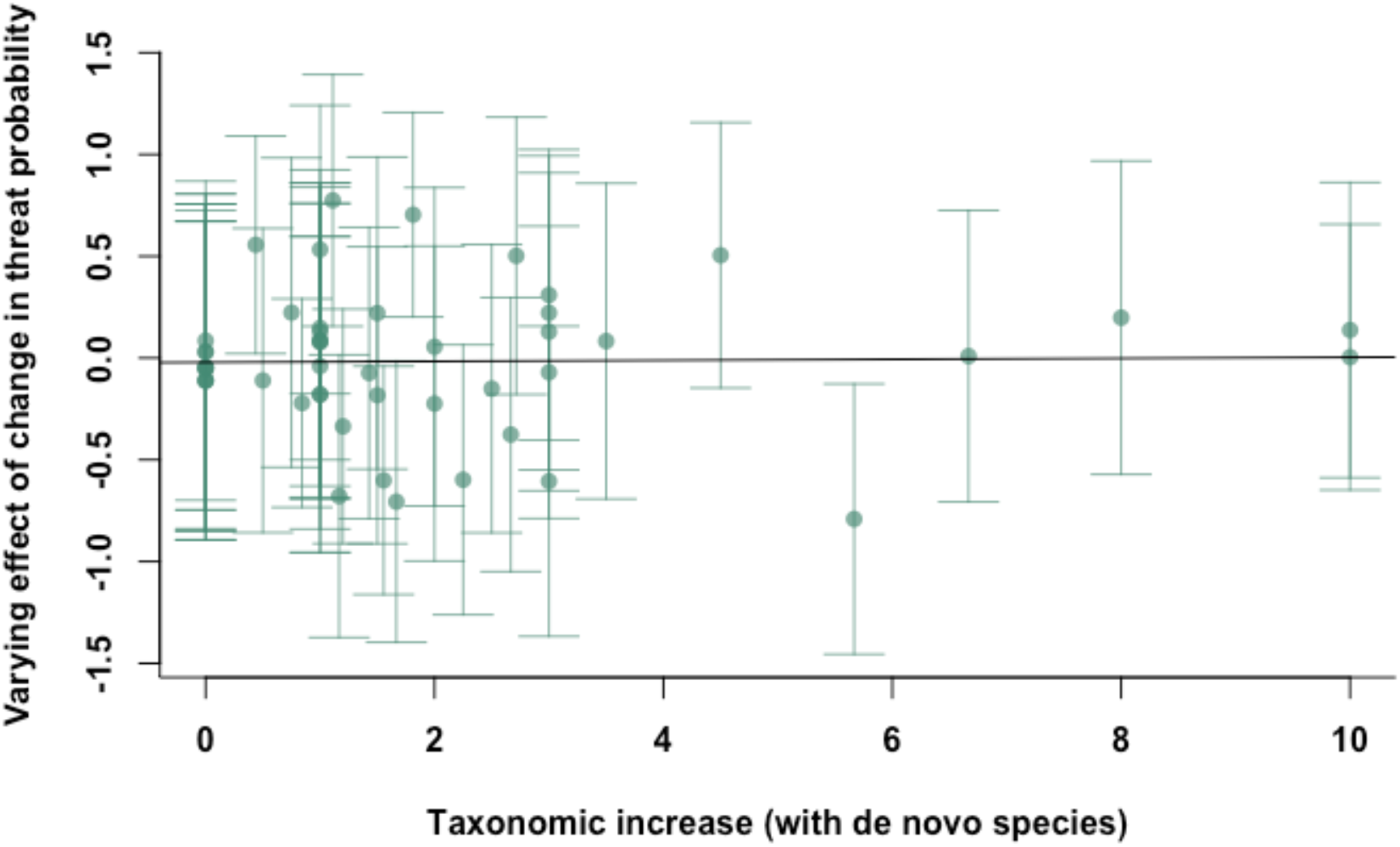
Taxonomic increase including *de novo* species descriptions versus varying effect of change in threat probability (i.e., our measure of genus level change in threat probability between 1982 and 2016; see Methods for an explanation of how this was estimated) (β= 0.001; p= 0.975; Table S7). Vertical bars indicate standard errors on varying effect of change in threat probability values.

## DISCUSSION

Our results support neither a strong biological nor a strong interest-driven mechanism for splitting across primate genera. If the naming of new species under new taxonomic approaches captures incipient speciation, we predicted that diversification rate should predict taxonomic increases. However, diversification rate was not a significant predictor of taxonomic increases in our models. We also tested the prediction that splitting is motivated by conservation interest (Karl & Bowen, 1999; Isaac *et al*., 2004) or by increasing molecular research within certain taxa (Zachos *et al*., 2013; Zachos & Lovari, 2013). However, we found no evidence that research effort in either of these areas was associated with the amount of taxonomic splitting observed across genera. We did find that initial genus size predicts increased splitting, but this effect decelerates as genera approach the largest sizes. This could indicate that splitting is being driven by variables not captured by our models. For example, genera that already contained many species in 1982 may have been more likely to have closely related populations described as separate species under the BSC (e.g., if we knew more about their hybrid statuses compared to other clades). In this case, we may have already discovered much diversity in these large clades prior to applications of the PSC and new molecular techniques. Future studies could aim to elucidate the origins of this decelerating association, however, our results indicate it is not explained by the rate at which lineages diverge (which should capture the presence of cryptic species) or research effort in the two fields studied here.

While inspection of the raw data suggested that genus level measures of splitting, threat probability, and change in threat probability all varied by region, we found that there was no cohesive signal of taxonomic splitting leading to higher threat probability across these genera and regions. Therefore, while there are idiosyncratic regional differences in both splitting and threat probability there was no evidence for causal links between the two. We do note that while the act of splitting itself does not seem to be having an overwhelming effect on threat status, it is still possible splitting may have uncaptured negative consequences downstream in conservation management (e.g., loss of genetic variation in captive breeding programs; Zachos, 2013). However, our analysis sends a positive message that splitting is not currently a significant determinant of the relative conservation priority of primate clades, consistent with recent findings in birds (Simkins *et al*., 2020).

Importantly, we note that our analyses have some limitations. First, quantifying research effort of any kind is difficult due to the abundance of work published in different media. As such, our estimates of research effort in conservation and molecular genetics may overlook some types of research. More work could be done to determine if additional estimates of research bias can explain increases in species numbers. For instance, cumulative funding estimates from various sources (e.g., the IUCN, non-governmental organizations and regional governments) per taxon could provide an additional or alternative measure of conservation interest. Second, it was necessary to remove eight non-monophyletic genera (some of which have undergone substantial splitting) from analyses that included diversification rate, leading to a considerable reduction in statistical power for those analyses. Third, due to changes in Red List criteria, species status, and lack of information about which species had been assessed in the 1980s, we were not able to consider differences in threat status severity (e.g., Vulnerable versus Endangered) when considering associations between taxonomic splitting and changes in threat score through time. Future studies could look at the association between splitting and changes in weighted measures of threat score (e.g., RLI) over time using a more recent starting point (i.e., after 1982) where Red List criteria become stable and there is available information on which non-threatened species have been evaluated.

Overall, we do not find support for biological processes or research bias driving taxonomic splitting across primate genera. We also find no cohesive signal of splitting leading to higher threat probabilities. Generally speaking, relying on species as the central unit of conservation and primary object of biological study behooves taxonomy to remain stable, while changing ideas about the concept of species makes taxonomy inherently unstable (Mayr, 1996). We suggest that areas of research requiring consistent estimates of diversity (e.g., conservation, macroecology, or evolutionary biology) may benefit from (i) weighing evolutionary distinctiveness when determining how species are listed/treated if attempting to capture true biological diversity (see, e.g., Redding & Mooers, 2010; Redding *et al*., 2015); or, for applied conservation specifically (ii) shifting more resources toward regional management efforts that are less likely to be influenced by changing species designations. It is well-known that closely related species are more similar to one another than they are to more distantly related taxa. Thus, treating all species independently and of equal weights in conservation listing may not lead to desired outcomes (Redding & Mooers, 2010). As of 2016, approximately 60 percent of all primate species were threatened with extinction according to the IUCN Red List of Threatened Species (Estrada *et al*., 2017), making it imperative that conservation efforts are spent wisely to ensure optimal conservation of primate biodiversity writ large.

## ACKNOWLEDGEMENTS

This work was supported by the Natural Sciences and Engineering Research Council of Canada (NSERC) Discovery Grants Program (SMR and AOM, grants #2017-04720 and #2019-04950) and the Canada Foundation for Innovation (CFI) (SMR, grant #29433). MJAC was supported by awards from the Biodiversity, Ecosystem Services, and Sustainability (BESS) program and McGill University. We thank the Reader and Guigueno labs at McGill, the Crawford Lab for Evolutionary Studies at SFU, Brian Leung, Hans Larsson, William Wcislo, and Isabella Capellini for feedback on portions of the manuscript and study design. Special thanks to Aiman Hadif for his assistance in collecting and organizing data and to Paul Sims and Dan Greenberg for statistical advice. We also thank Sandro Lovari, Frank Zachos, and one anonymous reviewer for their useful comments on earlier drafts.

## DECLARATIONS OF INTEREST

None.

Data and code accompanying the manuscript are available on Zenodo: https://doi.org/10.5281/zenodo.6614874

## Notes

### Competing Interest Statement

The authors have declared no competing interest.

### Summary of Updates

This updated version of the manuscript focuses on the causes and consequences of taxonomic splitting, and removes emphasis from the comparison of species concepts.

