## Supplementary Materials for "Predictors of taxonomic splitting and its role in primate conservation"

Maria J.A. Creighton[a], Alice Q. Luo[a], Simon M. Reader[a] & Arne Ø. Mooers[b]
[a]Department of Biology, McGill University, Montreal, Quebec, Canada
[b]Department of Biological Sciences, Simon Fraser University, Burnaby, British Columbia, Canada

### SUPPLEMENTARY MATERIALS AND METHODS:

#### *Data*

*Research Effort*

Estimates of research effort for each genus listed by Honacki *et al.* (1982) in the fields of conservation and molecular genetics were determined through an extensive literature review of papers in the Web of Science Core Collection (Tables S1, S2 and S3). The search included studies published between 1983 – the year the ‘phylogenetic species concept’ (PSC) was first proposed, and 2016 – the year the IUCN species list and associated data documented in Estrada *et al.* (2017) were collected. In this study we used the 50 genera listed in Honacki *et al.* (1982), many of which have since been further separated into multiple genera. Thus, when appropriate we included new genera names in the literature search in addition to those listed by Honacki *et al.* (1982) to ensure our search returned all relevant papers (Table S3). Four genera included names that were associated with unrelated topics or doubled as a common name for species in other genera (e.g., genus *Lemur*). For these genera, all possible species names were added to the search to prevent irrelevant hits (Table S3).

To obtain papers in the field of conservation we searched the genus name/names AND “conservation”. To obtain papers in the field of molecular genetics we searched the genus name/names AND the following terms: "mitochondrial DNA" OR "barcoding" OR "bar-coding" OR "cytochrome b" OR "phylogeography" OR "microsatellites" OR "micro-satellites" OR "population genetics" OR (species AND genetics) OR (taxonomy AND genetics) OR "SNP". All hits from these initial searches were reviewed on a case-by-case basis and NOT terms were created to eliminate unrelated hits (see Tables S1 and S2). We did this rather than manually removing records from the research effort so that searches could be replicated. We reran searches on a subset of genera to confirm that this procedure did not eliminate relevant papers. Searches for some genera returned hits that were not relevant to the genus of interest but were relevant to others in which case NOT terms specific to each genus were included in the search (see Table S3). When it came to molecular genetics, we were interested in the contribution of new molecular work that could lead to splitting, therefore, results in this search were filtered by articles and letters to exclude material like reviews that would not make such contributions. Once the final search terms were established, the searches were repeated with the full list of NOT terms and the number of results returned for each genus using these terms was recorded, providing us with a comprehensive estimate of research effort on each topic per genus. These searches can be replicated in the Web of Science databases using the terms included in Tables S1, S2 and S3.

Table S1: Terms used for literature review to obtain an estimated research effort in the field of conservation per genera in Honacki *et al.* (1982) between 1983 and 2016. Terms in inverted commas are those entered into the search engine. "Genus A", Genus B" etc. represent the alternative genus names presented in Table S3. Asterisk in search term denotes a wildcard character representing any group of characters, including no character.

| Topic | Search |
|---|---|
| Conservation | **Database: Web of Science Core Collection**<br>**Year Range: 1983-2016**<br><br>*"GenusA" OR "GenusB"* **[Search field: TOPIC]**<br><br>**AND** "Conservation" **[Search field: TOPIC]**<br><br>**NOT** "Pathalog*" OR "Seminal" OR "Semen" OR "Protein" OR "Numerical Competence" OR "*sterone" OR "Memory" OR "Chromosome" OR "DNA" OR "Molecul*" OR "Euterpe edulis" OR "Mico-particles" OR "Gut passage time" OR "MHC" OR "PAXBP1" OR "Cathemerality" OR "Arabic" OR "Vomeronasal Receptor" OR "CT Repeats" OR "Butterfly" OR "microRNA" OR "Mirrors" OR "Kinematics" OR "geogenetic" OR "archaeology" OR "monuments" OR "quantum" OR "political crisis" OR "neutral theory" OR "baobabs" OR "tor macrolepis" OR eichhornia crassipes" or "ethnobotany" OR "movement corridors" OR "null errors" OR "skeletal indices" OR "resource defense" OR "energetic demand" OR "chromosomes" OR "floristic composition" OR "sickness behavior" OR "appearance-reality" OR "object permanence" OR "number processing" OR "quantity judgement" OR "quantity discrimination" OR "visual illusions" OR "oil extraction" OR "object manipulation" OR "piagetian" OR "personality" OR "forest regeneration" OR "Bergmann's Rule" OR "female dominance" OR "publication" OR "rhinoceros" OR "photopigments" OR "opsin" OR "FSH" OR "IFITM10" OR "IFITM5" OR "energy expenditure" OR "myopia" OR "fovea" OR "museum" OR "cacao cultivation" OR "cabruca plantation" OR cabrucas" OR "discrimination learning" OR "architecture" OR "tocoplasma goondii" OR "dung beetle" OR "meat science" OR "year span" OR "los toxtlas" OR "lipid droplets" OR "nephropathy" OR "battelle" OR "luncheon address" OR "fruit-eating fish" OR "parrots" OR "nest fate" OR "bird community" OR "retinal development" OR "edinger-westphal nucleus" OR "fur mites" OR "alliances" OR "energy metabolism" OR "nucleotide sequence" OR "transcriptional effector" OR "lymphotropic" OR "RNA secondary" OR "ischemic stroke" OR "lifetime achievement award" OR "number representation" OR "metabolic activity" OR "natureculture" OR "fruit fly" OR "pluripotent stem cells" OR "locus ERVWE1" OR "duffy blood group system" OR "p58" OR "coraco-clavicular joint" OR "kaya" OR "Shannon-Weiner index" OR "kisspeptin" OR "linkage map" OR "killer whales" OR "female athlete" OR "acetyl salicylic acid" OR "periaqueductal gray" OR "DQA1" OR "dentate granule cells" OR "descartes" OR "haplotype network analysis" OR "OCOM-6" OR "Participatory risk mapping" OR "Fribourg-Blanc genome" OR "browser" OR "atherosclerosis" OR "rapid lateral flow" OR "VBEKAP" OR "super-eruption" OR "fossil mammals" OR "evolution of dance" OR "biate tribe" OR "antiphonal songs" OR "energy conservation" OR "strongyle egg excretion" OR "divergence date" OR "phylogenic niche conservation" OR "spatial autocorrelation" OR "conservation of heat" OR "local terrestrial tradition" OR "first fossil record" OR "nonhuman primate amygdala" OR "skin cell proliferative potential" OR "nuclear hormone receptors" OR "prehistoric populations" OR "last glacial maximum" OR "prehistoric demographic event" OR "sugar-rich fruit pulp" OR "quantity conservation"<br>**[Search field: TOPIC]**<br><br>**AND** **Insert 'Species Names (if applicable)' – see Table S3** **[Search field: TOPIC]**<br><br>**NOT** **Insert 'Genus-specific NOT Terms (if applicable)' – see Table S3** **[Search field: TOPIC]** |

<u>Table S2:</u> Terms used for literature review to obtain an estimated research effort in the field of molecular genetics per genera in Honacki *et al.* (1982) between 1983 and 2016. Terms in inverted commas are those entered into the search engine. “Genus A”, Genus B” etc. represent the alternative genus names presented in Table S3. Asterisk in search term denotes a wildcard character representing any group of characters, including no character.

| Topic | Search |
|---|---|
| Molecular genetics | **Database: Web of Science Core Collection**<br>**Year Range: 1983-2016**<br><br>"*GenusA*" OR "*GenusB*" **[Search field: TOPIC]**<br><br>**AND** "Mitochondrial DNA" OR "Barcoding" OR "Bar-coding" OR "cytochrome b" OR "Phylogeography" OR "Microsatellites" OR "Micro-satellites" OR "Population Genetics" OR (Species AND Genetics) OR (Taxonomy AND Genetics) OR "SNP" **[Search field: TOPIC]**<br><br>**NOT** "Disease" OR "Virus" OR "Parasite" OR "Single Origin" OR "Color Vision" OR "Colour Vision" OR "Cyprinine fishes" OR "Schizothorax" OR "Ground boa*" OR "Tenrecidae" OR "Climate-Change" OR "Galaxy Tools" OR "Primatology in China" OR "Hand morphology" OR "Human Genome" OR "Bifidobacteri*" OR "Plasmodium" OR "Spondias" OR "Vestigial structures" OR "Social behavior, reproductive strategies" OR "Tuberculosis" OR "Social dynamics of male" OR "Skull shape" OR "Diet diversity" OR "Functional cues" OR "Video stimuli" OR "Human specific" OR "Carrion fly" OR "Arctocephalus" OR "Bushmeat" OR "Elephants" OR "Human-Population Genetics" OR "Antimicrobial" OR "Vertebral formulae" OR "HIV-1*" OR "Genotoxicity" OR "Habitat fragmentation" OR "Primer design" OR "Plant DNA" OR "Parasitic lice" OR "Primates of Gashaka" OR "Marmot" OR "Captive breeding" OR "Captive-bred" OR "Behavioral genetics" OR "Toxicolo*" OR "Pharmacolo*" OR "Blood type" OR "Tissue expression" OR "Behavioral plasticity" OR "Medicine" OR "Immuno*" OR "Infection" OR "Reproductive efficiency" OR "Pinworm" OR "Diffusion tensor" OR "Female homosexual behavior" OR "Cryodamage" OR "Adult mortality" OR "iSCNT" OR "Resources for genetic management" OR "Hearing loss" OR "Macular degeneration" OR "Biomedical Information Research Network" OR "Korean cattle" OR "Sinus volume" OR " Alcoholism" OR "Dominance rank" OR "Therapeutic cloning" OR "Literature survey" OR "Grasshopper*" OR "Leptin" OR "Macaca-rabbit" OR "Nitrate tolerance" OR "Human evolution" OR "South American rodents" OR "hepatic CYP2C" OR "Baroreceptor-related neurons" OR "Nasalis posterior" OR "Salamander" OR "Gasterophilus nasalis" OR "Grooming bouts" OR "Rainfall" OR "Craniofacial" OR "Diabetes" OR "Sexual size dimorphism" OR "Mesopithecus" OR "Hybrid origin of the kipunji" OR "Schistosoma mansoni" OR "Limb bones" OR "Acinonyx jubatus" OR "Early life mortality" OR "Human pigmentation" OR "Modern Humans" OR "Balsaminaceae" OR "Meat consumption" OR "Tracking of a gorilla death" OR "Antrocaryon klaineanum" OR "ATP synthetase in orangutan" OR "Human mitochondrial DNA" OR "Maximal lifespan" OR "Mummified baboon" OR "Kin bonds" OR "Kinship" OR "Bipolar disorder" OR "Cultural differences" OR "Sex-specific dispersal" OR "Nest-building" OR "Vocalizations" OR "Neuroticism" OR "Stone handling" OR "Bowhead whale" OR "Canis rufus" OR "Caledonian crows" OR "Paternity" OR "Black rhinoceros" OR "Eptesicus fuscus" OR "4q syndrome" OR "Monozygotic twin" OR "Autism spectrum" OR "Skeletal analysis" OR "Furcifer lateralis" OR "Long-fingered bats" OR "Jomon period" OR "Cranial nerve perforations"<br>**[Search field: TOPIC]**<br><br>**NOT** "Toxicolo*" OR "Pharmacolo*" OR "Immuno*" OR "Medicine" **[Search field: PUBLICATION NAME]**<br><br>**AND** **Insert 'Species Names (if applicable)' – see Table S3** **[Search field: TOPIC]**<br><br>**NOT** **Insert 'Genus-specific NOT Terms (if applicable)' – see Table S3** **[Search field: TOPIC]**<br><br>**Refined by: Article or Letter** |

Table S3: Genera names included in each literature review for research effort with species names and additional NOT terms included in each search where applicable. Genus names were matched to Honacki *et al.* (1982) from the IUCN species list documented in Estrada *et al.* (2017). Additional alternative genus names were obtained from the IUCN Red List of Threatened Species (2019). Asterisk in search term denotes a wildcard character representing any group of characters, including no character.

| **Genus Name/Names** | **AND Species Names** *(if applicable)* | **Molecular genetics genus-specific NOT Terms** *(if applicable)* | **Conservation genus-specific NOT Terms** *(if applicable)* |
|---|---|---|---|
| **"Allocebus"** | | | |
| **"Cheirogaleus"** | | | |
| **"Microcebus" OR "Mirza"** | | "Saguinus mystax" OR "Sportive lemurs (Lepilemur, Primates)" OR "Bornean orang-utans (Pongo pygmaeus)" OR "Black-and-white ruffed lemur" | |
| **"Phaner"** | | | |
| **"Hapalemur" OR "Prolemur"** | | | |
| **"Lemur" OR "Eulemur"** | "albifrons" OR "cinereiceps" OR "collaris" OR "coronatus" OR "fulvus" OR "macaco" OR "flavifrons" OR "mongoz" OR "rubriventer" OR "rufus" OR "sanfordi" OR "sanfordi" OR "albocollaris" OR "mayottensis" OR "petterus" OR "catta" OR "rufifrons" | "Microcebus" OR "Mouse lemur" | "divergence from other mouse lemur clades" OR "preferred habitat type of Prolemur simus" OR "the Critically Endangered greater bamboo lemur Prolemur simus" OR "(Tarsius spectrum) in Tangkoko Nature Reserve" OR "presence of Prolemur simus at 18 sites" OR " critically endangered greater bamboo lemur Prolemur simus" OR "(Allocebus trichotis) to determine habitat needs" OR "the third species, Microcebus mittermeieri" OR "the third species, Microcebus mittermeieri" OR "205 Microcebus ravelobensis" OR "focal individuals of the weasel sportive lemur" OR "low reproduction rate for Lepilemur edwardsi" OR "Avahi occidentalis extends north and east of the Betsiboka River" OR "genetic structure of the solitary grey mouse lemur" OR "physiological parameters in healthy wild Varecia populations" OR "uneven distribution pattern of the golden-brown mouse lemur" OR "subspecies of the single species Varecia variegata" OR "First discovery of the hairy-eared dwarf lemur (Allocebus trichotis)" OR " almost extinct lemur species, Allocebus trichotis" OR "large breeding center for ruffed lemurs (Varecia variegata)" OR "Rediscovery of Allocebus-Trichotis Gunther" |

*(Continues on next page)*

| | | | |
|---|---|---|---|
| **"Varecia"** | | "Mouse lemur" | "Cheirogaleus" OR "Lemur" OR "Canarium" OR "chimpanzee" |
| **"Lepilemur"** | | | |
| **"Indri"** | | | |
| **"Avahi or Lichanotus"** | | | |
| **"Propithecus"** | | "Lemur catta" | "Ateles" OR "Chiropotes" |
| **"Daubentonia"** | | | |
| **"Arctocebus"** | | | |
| **"Loris"** | "tardigradus" OR "lydekkerianus" | | "Mirza" OR "Nycticebus" |
| **"Nycticebus"** | | | |
| **"Perodicticus"** | | | |
| **"Galago" OR "Galagoides" OR "Paragalago" OR "Sciuricheirus" OR "Euoticus"** | "elegantulus" OR "pallidus" OR "demidoff" OR "thomasi" OR "orinus" OR "rondoensis" OR "granti" OR "cocos" OR "zanzibaricus" OR "senegalensis" OR "gallarum" OR "moholi" OR "matschiei" OR "alleni" OR "alleni cameronensis" OR "gabonensis" OR "makandensis" OR "demidovii" OR "inustus" OR "tonsor" OR "apicalis" OR "talboti" OR "pallida" OR "demidovii" OR "udzungwensis" OR "zanzibaricus" OR "nyasae" OR "bradfieldi" OR "cameronensis" | | |
| **"Otolemur"** | | | |

*(Continues on next page)*

| | | |
|---|---|---|
| **"Tarsius" OR "Carlito" OR "Cephalopachus"** | | |
| **"Callimico"** | "Owl monkey*" | |
| **"Cebuella"** | | "Alouatta" |
| **"Callithrix" OR "Mico" OR "Calibella"** | "Owl monkey*" | "Saguinus" OR "Lemur" OR "litter size of wild cotton-top tamarins" OR "reproductive potential of wild- and captive-born golden-headed lion tamarins" OR "two tamarin species (Saguinus fuscicollis and S. oedipus)" |
| **"Leontopithecus"** | | "Saguinus" OR "threats to populations of Alouatta guariba clamitans " |
| **"Saguinus" OR "Leontocebus"** | "Leontopithecus caissara" OR "Rondon's Marmoset" OR "Cebus apella paraguayanus" OR "skull of Catarrhini" OR "masticatory apparatus of Galagos" OR "cranial diversification of neotropical monkeys" | "cuxius formed associations" OR "breast height, 42.2 +/- 21.9 cm" OR "(Callicebus cupreus) to adapt to forest edges" OR "Callimico goeldii is a rare primate" OR "dung in latrines" OR "matrix diagonalization" OR "Sleeping Sites of Rhinopithecus brelichi" OR "Leontopithecus rosalia Linnaeus, 1766" OR "night monkeys Aotus vociferans" OR "Habitat use by Chiropotes satanas utahicki " OR "20 Alouatta species and subspecies" |
| **"Alouatta"** | "Owl monkey*" OR "Hooded capuchin" OR "Squirrel monkey" | "Rhinopithecus" OR "Lemur" OR "Harpia" OR "Coleoptera" OR "Eulemur" OR "Scarabaeidae" OR "Chiropotes" OR "Hylobates" OR "population of white-bellied spider monkeys (Ateles belzebuth)" OR "1 adult and subadult brown spider monkeys " OR "ability of spider (Ateles geoffroyi)" OR "Ateles marginatus, is endemic to Brazilian Amazon" OR "A. geoffroyi live in highly fragmented landscapes" |
| **"Aotus"** | "Woolly monkey" OR "Saguinus" OR "MHC Polymorphisms" | |
| **"Ateles"** | | "Rhinopithecus" OR "Papio" OR "Cebidae" OR "Inga" OR "(Alouatta palliates, Alouatta pigra and" OR "Demographic features of Alouatta pigra populations" OR "20 Alouatta species and subspecies" |

*(Continues on next page)*

| | | |
|---|---|---|
| **"Brachyteles"** | "Howler monkey*" | "Chiropotes" OR "Tapir" OR "brown howler monkey (Alouatta clamitans)" OR "brown-howler-monkey Alouatta guariba clamitans " OR "20 Alouatta species and subspecies" |
| **"Cacajao"** | | "Pitheciid" |
| **"Callicebus" OR "Cheracebus" OR "Plecturocebus"** | | "Pitheciid" |
| **"Cebus" OR "Sapajus"** | "Saguinus mystax" OR "Howler monkey*" | "Papio" OR "calithrix" OR "Presbytis" OR "Harpia" OR "Dasypus" OR "Leopardus" OR "Macaca" OR "9 healthy captive ring-tailed lemurs" OR "cuxius (genus Chiropotes) form" OR "northern bearded sakis (Chiropotes sagulatus) in Guyana" OR "recently-discovered titi, Callicebus coimbrai" OR "last in-depth review of Saimiri biology" OR "L. rosalia is a legitimate seed disperser" OR "endemic Chiropotes satanas utahicki" OR "20 Alouatta species and subspecies" OR "Central American squirrel monkeys, Saimiri oerstedi" |
| **"Chiropotes"** | | "Pitheciid" OR "Pitheciinae" |
| **"Lagothrix" OR "Oreonax"** | | "Nomascus" OR "Burseraceae" OR "Varecia" OR "Brachyteles" OR "Chiropotes" |
| **"Pithecia"** | | "20 Alouatta species and subspecies" |
| **"Saimiri"** | "Colobine monkeys" OR "Saguinus" OR "Owl monkey*" OR "Howler monkey*" | "rhesus monkeys (Macaca mulatta) succeeded" OR " 20 Alouatta species and subspecies" |
| **"Allenopithecus"** | | |
| **"Cercocebus" OR "Lophocebus"** | "Rungwecebus" | "drill population ecology" |
| **"Cercopithecus" OR "Allochrocebus" OR "Chlorocebus" OR "Miopithecus"** | | "Alouattta" OR "Macaca" OR "Canis" OR "Erythrocebus" OR "endemic Udzungwa red colobus (Procolobus gordonorum)" OR "drills on 25 occasions" OR "conflict in the guanaco (Lama guanicoe)" OR "15,000 Papio hamadryas hamadryas" OR "gorilla nest-site densities" OR "leopard was ascertained only " |

*(Continues on next page)*

| | | |
|---|---|---|
| **"Erythrocebus"** | | |
| **"Colobus" OR "Procolobus" OR "Piliocolobus"** | | "Chiropotes" OR "Eulemur" OR "Callicebus" OR "Presbytis" OR "Simias" OR "Semnopithecus" OR "Rhinopithecus" OR "Trachypithecus" OR "Lepilemur" OR "village-dwelling populations (Lagwa and Akpugoeze)" OR "cercopithecid conservation" OR "Supplementation in Black howlers (Alouatta pigra)" OR " Boutourlini's blue monkeys (Cercopithecus mitis boutourlinii)" OR "Foraging Behavior of Red Howler Monkeys" OR "drills on 25 occasions" OR "Cercocebus sanjei in the Udzungwa Mountains" |
| **"Macaca"** | "Rhinopithecus roxella*" OR "Rhinopithecus bieti" OR "Infra-order catarrhini" OR "Southern African baboons" OR "Hylobates lar" OR "Wild orangutans" OR "captive group of chimpanzees (Pan troglodytes)" OR "Cercopithecus aethiops aethiops" OR "wild living community of Bonobos (Pan paniscus)" OR "Bolivian squirrel monkeys (Saimiri boliviensis)" OR "Captive lowland gorillas and orangutans" OR "Population genetics in Eulemur" | "Rhinopithecus" OR "Neofolis" OR "Callithrix" OR "Cercopithecine" OR "Propithecus" OR "Cebus" OR "Theropithecus" OR "indian giant squirrel" OR "terrestriality in orangutans (Pongo spp.)" OR " zoo-housed Javan gibbons (Hylobates moloch)" OR "abituated orangutans (Pongo pygmaeus morio)" OR "crop-raiding by orangutans (Pongo abelii)" OR "discrimination skills of chimpanzees " OR "Panthera pardus have a catholic diet " OR "BFMS also contain C. vellerosus" |
| **"Nasalis" OR "Simias"** | | "Rhinopithecus" OR "Cercopithecidae" |
| **"Papio" OR "Mandrillus"** | | "Cercopithecine" OR "Prosopis" OR "West African savanna chimpanzees (Pan troglodytes verus)" OR "parasites of savanna chimpanzees (Pan troglodytes schweinfurthii)" OR "(Pan troglodytes) misperceived food" OR "Endangered Lion-Tailed Macaques (Macaca silenus)" OR "Cercocebus sanjei in the Udzungwa Mountains" OR "infecting greater spot-nosed monkeys (Cercopithecus nictitans)" OR "Tibetan macaques (Macaca thibetana) at Mt. Emei" |
| **"Presbytis" OR "Semnopithecus" OR "Trachypithecus"** | "Sichuan snub-nosed monkeys" | "Andrias" OR "Pongo" OR "Saimiri" OR "Pardofelis" OR "indian giant squirrel" OR "Simias concolor (simakobu or pig-tailed langur)" OR "(Rhinopithecus bieti) 14" OR "Food habits of tigers Panthera tigris" |

*(Continues on next page)*

| | | | |
|---|---|---|---|
| **"Pygathrix" OR "Rhinopithecus"** | | | "Endangered Shortridge's capped langur Trachypithecus shortridgei" OR "(Presbytis rubicunda) in Sabangau" OR "Responses of Cao Vit Gibbon (Nomascus Nasutus)" OR "(Trachypithecus delacouri) in Van Long Nature Reserve" |
| **"Theropithecus"** | | "Macaca arctoides" | |
| **"Hylobates" OR "Hoolock" OR "Nomacus" OR "Nomascus" OR "Symphalangus" OR "Brunopithecus"** | | "Cercopithecidae" OR "New World Monkey" | "Neofelis" OR "Saimiri" OR "Garcinia" OR "Choerospondias" OR "(Trachypithecus crepusculus) in the Wuliang Mountains" OR "one wild troop of northern pigtailed macaques " |
| **"Gorilla"** | | "Wild chimpanzees (Pan troglodytes)" OR "Unhabituated chimpanzees" OR "Leontopithecus caissara" OR "White-handed gibbon" OR "Common gibbon" | "Hylobates" OR "Alouatta" OR "Neofelis" OR "captive hamadryas baboons (Papio hamadryas hamadryas)" OR "Cercopithecus mitis kandt i, the golden monkey" |
| **"Pan"** | "paniscus" OR "troglodytes" | "natural populations of orang-utan (Pongo pygmaeus)" OR "Mitochondrial DNA diversity in gorillas" OR "Barbary macaques (Macaca sylvanus)" | "Leontopithecus" OR "Chlorocebus" OR "cercopithecids" OR "Japanese macaques (Macaca fuscata yakui) " OR "Vulnerable owl-faced monkey Cercopithecus hamlyni" OR "(Trachypithecus phayrei crepusculus) in Phu Khieo" OR " capuchin monkeys' quantification" |
| **"Pongo"** | | "Brown Spider Monkey" OR "Gorilla systematics" OR "Tooth morphology" OR "White-handed gibbon" | "Cebus" OR "Tapiridae" OR "Trypanosoma " OR "Nigerian/Cameroon chimpanzee (Pan troglodytes ellioti)" OR "chimpanzees' food preferences" OR "(Pan troglodytes) misperceived food" OR "Chimpanzees made judgments" OR "chimpanzees (Pan troglodytes) discriminate" |

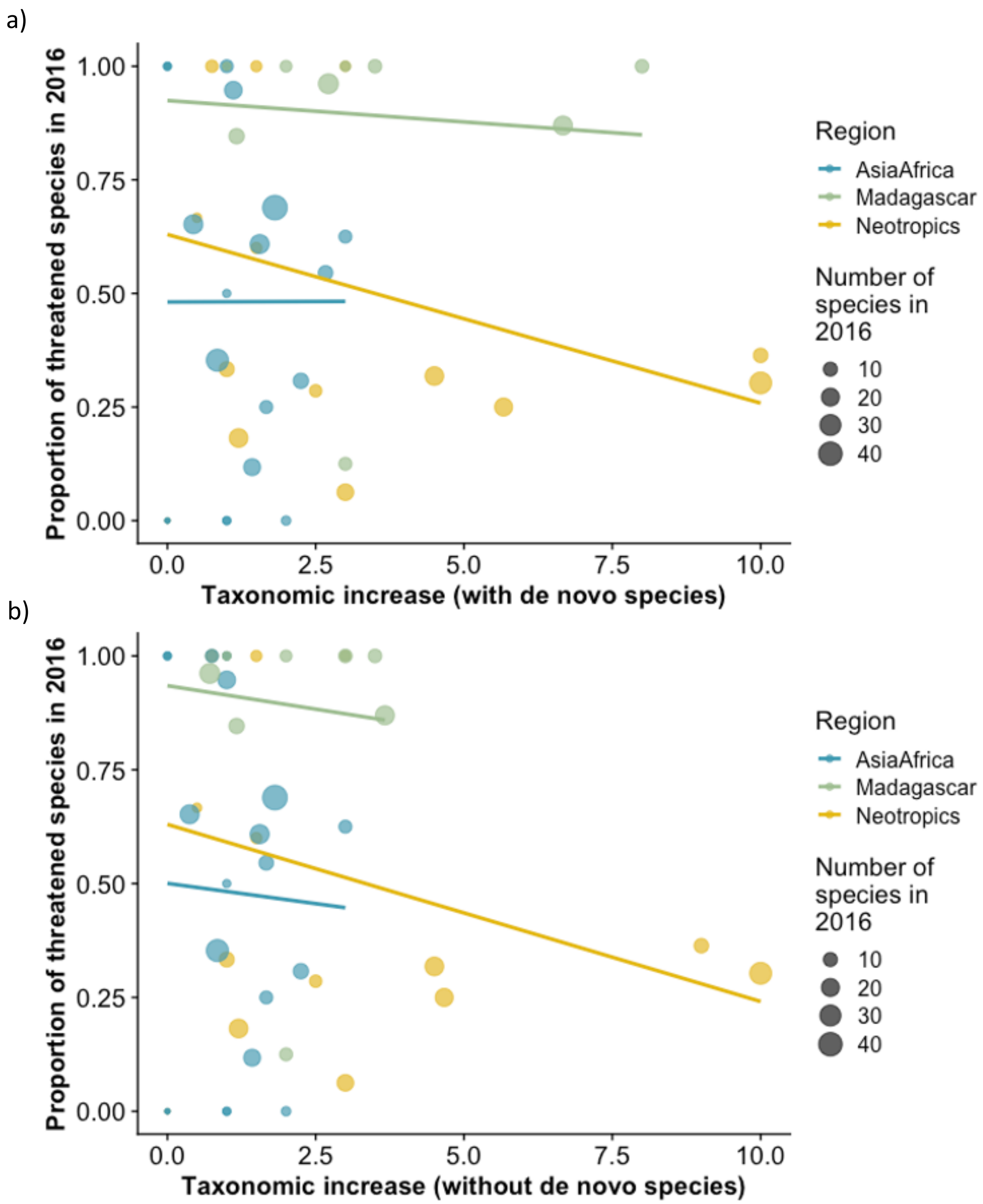


Figure S1: Scatterplot with trendlines showing the proportion of species identified as being threatened in primate genera in 2016 versus taxonomic increase a) including *de novo* species, and b) excluding *de novo* species, painted by region. Total number of species in each genus is indicated by point size.

a)

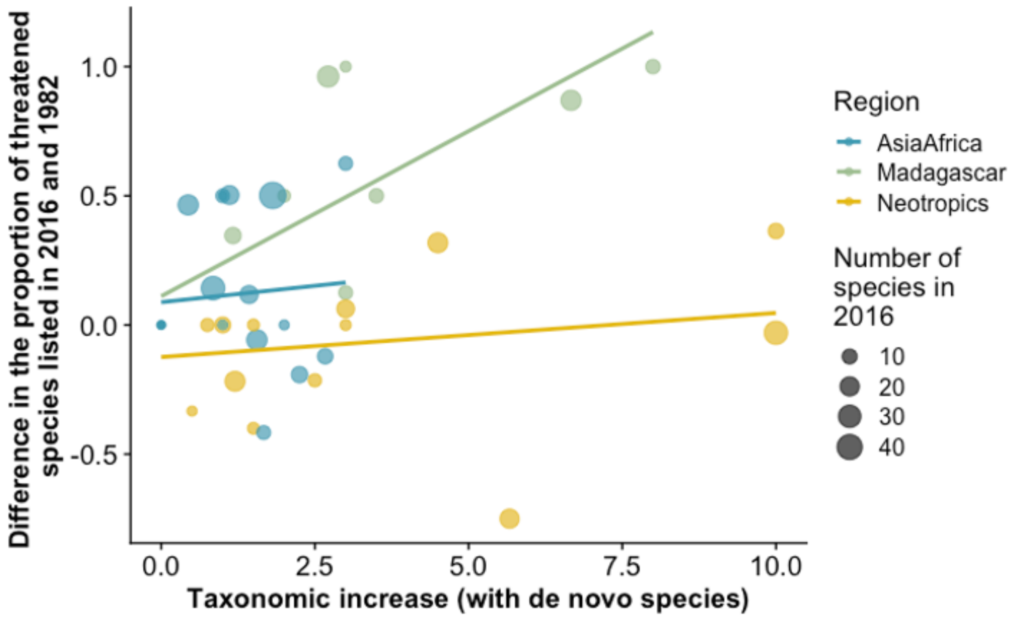


b)

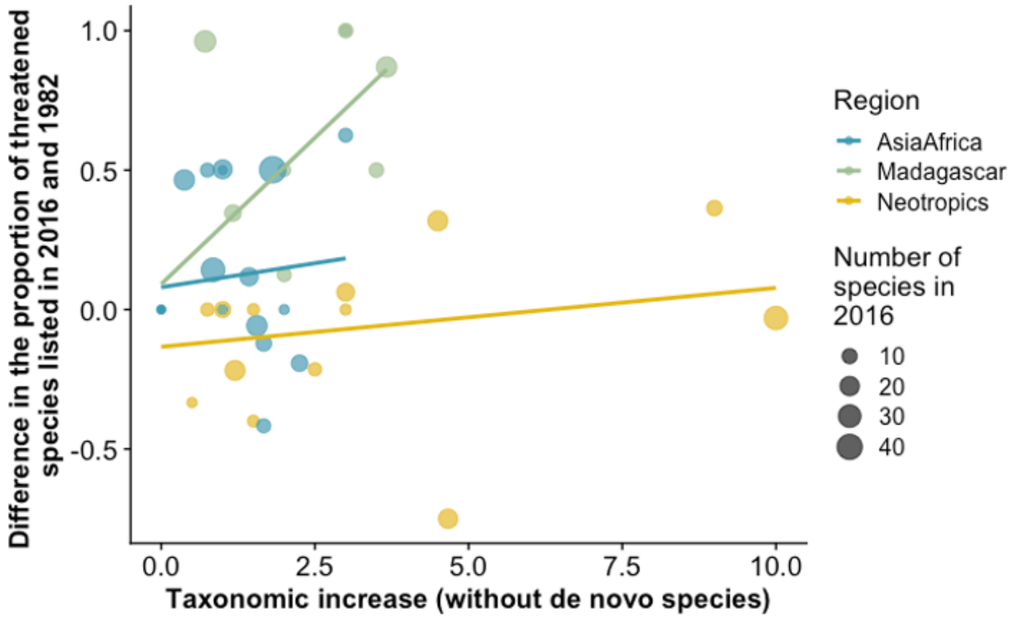


Figure S2: Scatterplot with trendlines showing the change in proportion of species identified as being threatened in primate genera between 2016 and 1982 (proportion at risk in 2016 – proportion at risk in 1982) versus taxonomic increase a) including *de novo* species, and b) excluding *de novo* species, painted by region. Total number of species in each genus is indicated by point size.

***Analysis***

*Model Choice*

Both taxonomic splitting and extinction risk are likely to be phylogenetically clustered, meaning that phylogenetic relationships are likely to account for some of the variation in splitting across genera. Phylogenetic models could be used to account for this influence of phylogeny, however, we used the genera listed by Honacki *et al.* (1982) as our level of observation. These genera are very conservative and do not always match up with recently described phylogenetic patterns. This means that there are several instances where genera described in Honacki *et al.* (1982) are non-monophyletic, making it unclear how to designate them a single branch in modern phylogenies. Notably, a majority of the phylogenetic clustering in taxonomic splitting is determined by differences in splitting rates among Madagascar, the Neotropics, and mainland Africa + Asia. Thus, we instead opted to use linear effects and mixed effects models where the effects of both region and family were considered to account for an influence of phylogeny, cognizant that some third-variable covariation may not be fully accounted for. After accounting for region, higher taxonomic designations (i.e., family) contributed little to no variance in any of our models indicating that phylogeny should not have significant influence on the results reported here.

To test whether taxonomic splitting over time is associated with a change in the proportion of threatened species within genera, we conducted a two-step analysis on species' threat probability between 1982 to 2016 using linear and linear mixed effects models. We note that a Bayesian framework could also be used to propagate error in this instance, however, to be consistent with models for our first question testing predictors of recent taxonomic increases we chose to use a frequentist approach that propagates error in a comparable way.

**SUPPLEMENTARY RESULTS:**

*Predictors of Taxonomic Splitting*

Table S4: Results of a generalized linear effects model testing the effect of conservation research effort on taxonomic splitting. Model set-up: number of species added to primate genera since 1982 (dependent variable) vs. original number of species in each genus in 1982 (included as a linear ($^1$) and quadratic ($^2$) term), conservation research effort and region using mainland Africa + Asia as the baseline (all independent variables), with observation ID included as a random effect. +P≤0.1; * P<0.05; ** P<0.01.

| **Effect** | ***With de novo species*** | | | | ***Without de novo species*** | | | |
|---|---|---|---|---|---|---|---|---|
| | Estimate | SE | z | p | Estimate | SE | z | p |
| (Intercept) | 0.904 | 0.220 | 4.107 | <0.001** | 0.873 | 0.214 | 4.084 | <0.001** |
| Conservation research effort | -0.220 | 0.164 | -1.342 | 0.180 | -0.157 | 0.156 | -1.011 | 0.312 |
| Region Madagascar | 0.581 | 0.338 | 1.721 | 0.085+ | 0.237 | 0.337 | 0.702 | 0.483 |
| Region Neotropics | 0.559 | 0.301 | 1.859 | 0.063+ | 0.597 | 0.288 | 2.071 | 0.038* |
| log(# of species in 1982)$^1$ | 7.647 | 1.057 | 7.234 | <0.001** | 7.178 | 1.017 | 7.060 | <0.001** |
| log(# of species in 1982)$^2$ | -1.529 | 0.849 | -1.802 | 0.072+ | -1.413 | 0.821 | -1.722 | 0.085+ |

Continuous variables scaled to have a mean of zero and standard deviation of one

Table S5: Results of a generalized linear effects model testing the effect of genetics research effort on taxonomic splitting. Model set up: the number of species added to primate genera since 1982 (dependent variable) vs. original number of species in each genus in 1982 (included as a linear ([1]) and quadratic ([2]) term), molecular genetics research effort and region using mainland Africa + Asia as the baseline (all independent variables) where observation ID included as a random effect. +P≤0.1; * P<0.05; ** P<0.01.

| **Effect** | ***With de novo species*** | | | | ***Without de novo species*** | | | |
|---|---|---|---|---|---|---|---|---|
| | Estimate | SE | z | p | Estimate | SE | z | p |
| (Intercept) | 0.889 | 0.222 | 4.012 | <0.001** | 0.867 | 0.213 | 4.074 | <0.001** |
| sqrt(Molecular genetics research effort) | -0.188 | 0.151 | -1.247 | 0.212 | -0.173 | 0.145 | -1.194 | 0.232 |
| Region Madagascar | 0.646 | 0.334 | 1.932 | 0.053+ | 0.281 | 0.330 | 0.851 | 0.395 |
| Region Neotropics | 0.550 | 0.305 | 1.806 | 0.071+ | 0.585 | 0.289 | 2.025 | 0.043* |
| log(# of species in 1982)[1] | 7.613 | 1.066 | 7.145 | <0.001** | 7.239 | 1.011 | 7.160 | <0.001** |
| log(# of species in 1982)[2] | -1.626 | 0.854 | -1.904 | 0.057+ | -1.449 | 0.818 | -1.772 | 0.076+ |

Continuous variables scaled to have a mean of zero and standard deviation of one

Table S6: Results of a generalized linear effects model testing the effect of diversification rate on taxonomic splitting. Model set up: the number of species added to primate genera since 1982 (dependent variable) vs. original number of species in each genus in 1982 (included as a linear ($^1$) and quadratic ($^2$) term), diversification rate and region using mainland Africa + Asia as the baseline (all independent variables) where observation ID included as a random effect. +P≤0.1; * P<0.05; ** P<0.01.

| **Effect** | ***With de novo species*** | | | | ***Without de novo species*** | | | |
|---|---|---|---|---|---|---|---|---|
| | Estimate | SE | z | p | Estimate | SE | z | p |
| (Intercept) | 0.623 | 0.288 | 2.166 | 0.030* | 0.603 | 0.274 | 2.204 | 0.028* |
| Diversification rate | 0.132 | 0.186 | 0.712 | 0.477 | 0.099 | 0.174 | 0.567 | 0.570 |
| Region Madagascar | 0.819 | 0.398 | 2.056 | 0.040* | 0.406 | 0.391 | 1.040 | 0.298 |
| Region Neotropics | 0.648 | 0.375 | 1.729 | 0.084+ | 0.708 | 0.352 | 2.011 | 0.044* |
| log(# of species in 1982)$^1$ | 5.303 | 1.186 | 4.469 | <0.001** | 4.877 | 1.126 | 4.330 | <0.001** |
| log(# of species in 1982)$^2$ | -1.890 | 0.931 | -2.030 | 0.042* | -1.830 | 0.887 | -2.064 | 0.039* |

Continuous variables scaled to have a mean of zero and standard deviation of one

*Taxonomic Splitting and Threat Score*

Table S7: Results of linear model comparing genus effect of change in threat probability (dependent) to increases in species numbers between 1982 and 2016. +P≤0.1; * P<0.05; ** P<0.01.

| **Effect** | ***With de novo species*** | | | | ***Without de novo species*** | | | |
|---|---|---|---|---|---|---|---|---|
| | Estimate | SE | t | p | Estimate | SE | t | p |
| (Intercept) | -0.014 | 0.069 | -0.196 | 0.845 | -0.007 | 0.069 | -0.101 | 0.920 |
| Taxonomic increase | 0.001 | 0.022 | 0.032 | 0.975 | -0.003 | 0.026 | -0.111 | 0.912 |

Continuous variables scaled to have a mean of zero and standard deviation of one

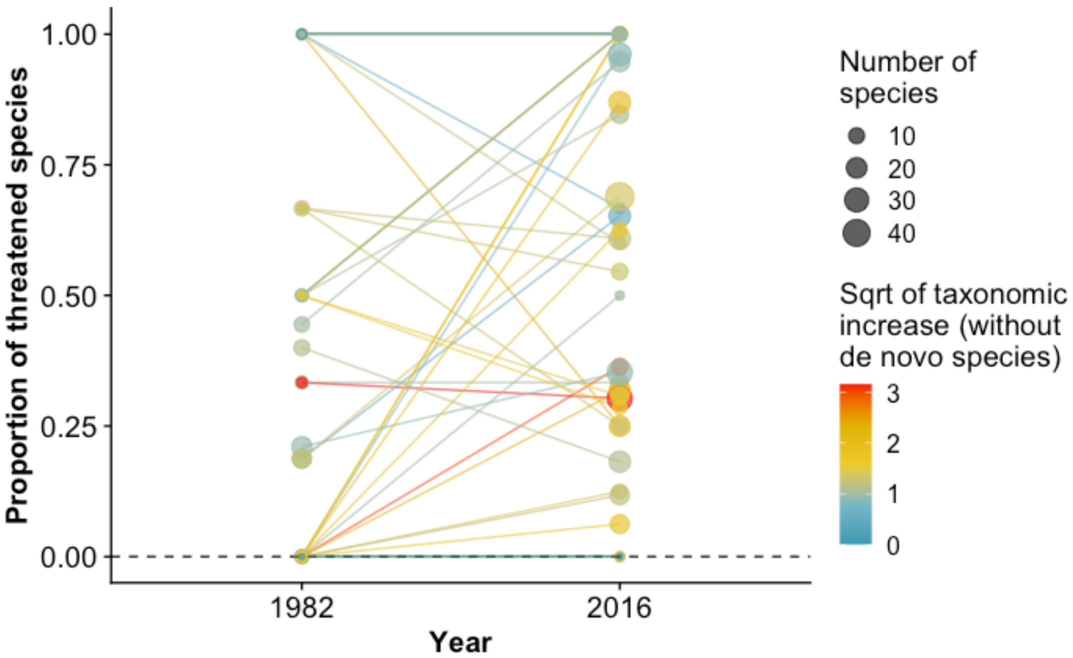


Figure S3: Scatterplots with trendlines showing the change in the proportion of species identified as being threatened in primate genera in 1982 and 2016 painted by the square root of taxonomic increase (excluding *de novo* species). Total number of species in each genus is indicated by point size.

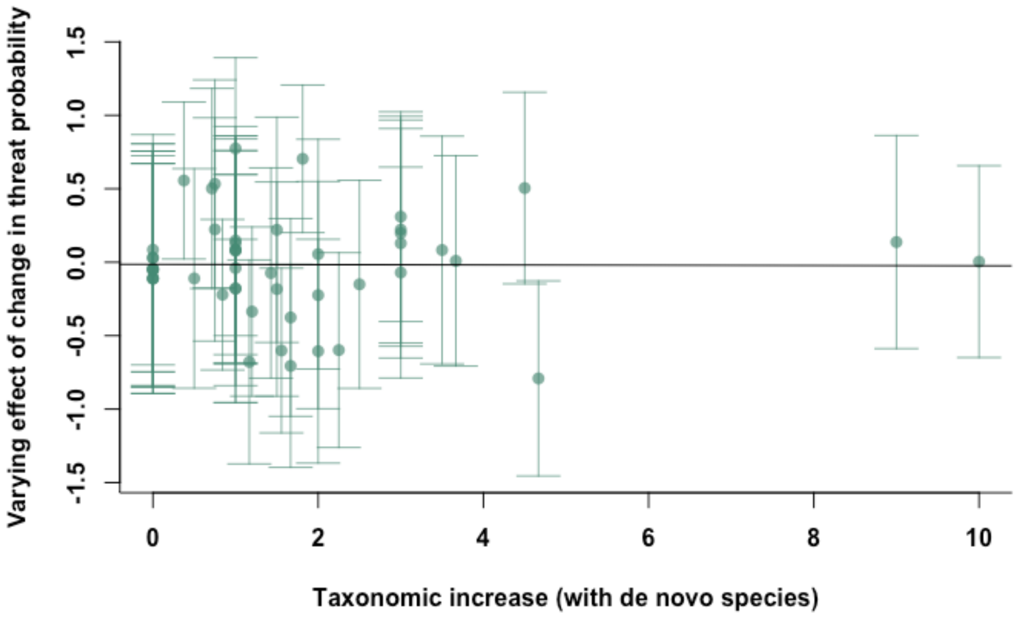


Figure S4: Taxonomic increase excluding *de novo* species descriptions versus varying effect of change in threat probability (i.e., our measure of genus level change in threat probability between 1982 and 2016; see Methods for an explanation of how this was estimated) (β= -0.003; p= 0.912; Table S7). Vertical bars indicate standard errors on varying effect of change in threat probability values.
